# STING causes replication stress and nascent DNA degradation via SAMHD1

**DOI:** 10.64898/2026.03.28.714577

**Authors:** Barbara Teodoro-Castro, Rafael Cancado de Faria, Elena V. Shashkova, Atika Malique, Madison B. Adolph, Lilian N.D. Silva, Susana Gonzalo

**Author notes:** Corresponding authors: Lilian N.D. Silva, PhD 1100 S. Grand Blvd St Louis, MO 63104, Susana Gonzalo, PhD 1100 S. Grand Blvd, St Louis, MO 63104. Equal contribution (Barbara Teodoro-Castro, Rafael Cancado de Faria).

## Abstract

STING is a key innate immune adaptor, classically activated by cytosolic DNA via cGAS–cGAMP to induce type I interferon signaling. While its cytoplasmic role is well defined, recent studies reveal that STING participates in non-canonical signaling pathways and localizes at the nuclear envelope and chromatin, where its functions remain poorly understood. In Hutchinson Gilford Progeria Syndrome (HGPS), a premature aging disease caused by expression of lamin A mutant protein named progerin, STING accumulates in the nucleus and drives chronic inflammation. Here, we show that replication stress (RS) is a trigger of STING nuclear accumulation and binding to chromatin. In addition, we uncover a previously unrecognized role for nuclear STING binding to nascent DNA and promoting RS in progeria and tumor cells. Mechanistically, STING contributes to replication fork slowing and stalling by limiting dNTPs availability. In addition, STING hinders replication fork protection/stability upon stalling, by facilitating MRE11-mediated nascent DNA degradation (NDD). We also find that STING contribution to depletion of dNTPs and NDD is mediated by SAMHD1. As such, SAMHD1 knockdown phenocopies STING abrogation in progeria cells and rescues replication fork speed and stability in STING-overexpressing tumor cells. These findings define a pathological STING–SAMHD1 axis that drives RS and genome instability in both progeria cells and tumor cells with elevated STING activity, uncovering a feedforward loop between innate immune signaling and impaired DNA replication.

**Highlights:**

- Replication stress in human fibroblasts triggers STING nuclear accumulation and an IFN response
- STING upregulation and nuclear accumulation hinders replication in progeria fibroblasts and U2OS tumor cells
- STING-induced replication stress features fork slowing/stalling and nascent DNA degradation
- STING-induced fork slowing/stalling is mediated by the dNTPase SAMHD1
- SAMHD1-enabled MRE11 activity is responsible for STING-induced nascent DNA degradation

## Introduction

The accumulation of self-DNA in the cytosol is a hallmark of aging and cancer, and a critical driver of inflammation and innate immune activation. The enzyme cGAS (cGAMP synthase) senses cytosolic DNA, and via the adaptor protein STING (STimulator of INterferon Genes) induces interferon (IFN) responses that promote anti-tumor immunity ^1–8^. However, chronic cGAS-STING activation drives age-related inflammation and senescence and contributes to diseases such as cancer, cardiovascular disease, and neurodegeneration ^9–11^.

The importance of the cGAS-STING signaling pathway has been shown in aging mice, in which blockade of STING suppresses inflammatory phenotypes of senescent cells, improves tissue function, and ameliorates functional decline ^12^. Our studies in a premature aging disease, Hutchinson Gilford Progeria Syndrome (HGPS), show that buildup of cytosolic DNA triggers an inflammatory cascade that includes an IFN response and a senescence-associated secretory phenotype (SASP) ^13,14^, which are ameliorated by depletion of cGAS or STING ^15^. Moreover, inhibiting STING reduces inflammation and cellular hallmarks of aging, ameliorates degeneration of adipose and vascular tissues, and extends lifespan of progeria mice (*Lmna^G609G/G609G^*) ^16^. These studies support a critical role for cGAS and STING in the inflammatory phenotypes of progeria and normal aging. Despite its therapeutic potential, the precise mechanisms whereby cGAS and STING impact cellular functions remain poorly defined.

STING resides in the ER and nuclear envelope (NE), becoming activated by cGAMP produced by cGAS upon cytosolic DNA recognition. Activated STING translocates to the Golgi/perinuclear compartment (PNC), where it activates TBK1, IRF3, and NFκB, inducing the expression of IFNs and pro-inflammatory cytokines that are released into the extracellular space. IFNs, via their receptors (IFNAR), trigger the JAK/STAT pathway and the expression of IFN-stimulated genes (ISGs) ^1,17,18^. We recently discovered a non-canonical cGAS-STING pathway driving sterile inflammation in aging, senescent, and progeria cells (from Hutchinson Gilford Progeria Syndrome patients) ^19^. Unlike canonical cGAS-STING signaling ^17,20–24^, this pathway proceeds without detectable increase in cGAMP production, STING trafficking to (PNC), or phosphorylation of STING and associated factors TBK1 and IRF3. Instead, it features increased nuclear STING, associating with the NE and chromatin. STING presence in the nucleus under resting conditions has been previously shown, and its binding to the inner nuclear membrane depends on lamin A^23,25^. Upon activation of the canonical cGAS-STING pathway in human primary dermal fibroblasts (HDF), the entire pool of STING is localized at the PNC, and the presence of STING at the nucleus appears lost ^19^. In contrast, progerin-expressing HDF show increased STING at the NE and in the nuclear interior in resting conditions and upon activation of the canonical cGAS-STING pathway with synthetic dsDNA. Importantly, this non-canonical pathway predominates in late passage ‘aged’ primary fibroblasts, and in senescent fibroblasts, coinciding with a reduced canonical responsiveness to foreign DNA ^19^. These findings reveal a previously unknown mechanism by which cellular aging blunts responses to foreign DNA while fueling key hallmarks of aging such as chronic inflammation. However, the causes of non-canonical cGAS-STING pathway activation and the consequences of STING accumulation in the nucleus are unknown.

Here, we show that the non-canonical cGAS-STING pathway in progeria cells features increased presence of STING in the nucleus, contributing to replication stress (RS). In tumor cells devoid of STING expression (U2OS cells), expression of STING causes RS. In both contexts, STING triggers replication fork slowing and stalling via depletion of dNTP pools, which is mediated by SAMHD1 (Sterile alpha motif and histidine-aspartic acid domain-containing protein 1), an interferon-stimulated gene (ISG) known to restrict viral replication by hydrolyzing dNTPs ^26^. In addition, STING causes replication fork deprotection and nascent DNA degradation (NDD) of stalled forks in progeria and tumor cells. Importantly, NDD is mediated by SAMHD1, which in addition to its dNTPase activity, functions facilitating the recruitment of the nuclease MRE11 to stalled replication forks ^27^. Our data reveals a toxic STING–SAMHD1 axis that drives RS and genome instability in both progeria cells and tumor cells with elevated STING activity.

## Results

### Replication stress drives nuclear accumulation of STING independent of canonical pathway activation

We previously showed that progerin expression alters STING localization, driving its accumulation in the nucleus and chromatin ^19^. This occurs in the absence of markers of classical cGAS-STING pathway activation such as elevated cGAMP production or STING phosphorylation and PNC trafficking ^19^. Given that progerin induces persistent replication stress (RS) ^13,28,29^, we asked whether RS itself could be sufficient to trigger this non-canonical STING behavior. To test this, we used hydroxyurea (HU), which depletes dNTPs and stalls replication forks, and single-stranded DNA (ssDNA) transfection, which mimics RS and it is known to activate DNA damage and immune response ^30^. These stimuli were compared to progerin expression using a doxycycline-inducible GFP-progerin system in human dermal fibroblasts (HDF-progerin), which recapitulates HGPS phenotypes within a few days of induction ^13,16^. Subcellular fractionation showed that HU, ssDNA transfection, and progerin expression (Doxy) lead to STING enrichment at the nuclear and chromatin-bound fractions, with progerin having the strongest effect (**Figure 1A**). Immunofluorescence revealed that HU and ssDNA transfection induced a distinct nuclear rim-like STING localization pattern (**Figure 1B**). This contrasted with the perinuclear accumulation seen after dsDNA transfection, which reflects canonical STING activation.

**Figure 1:**
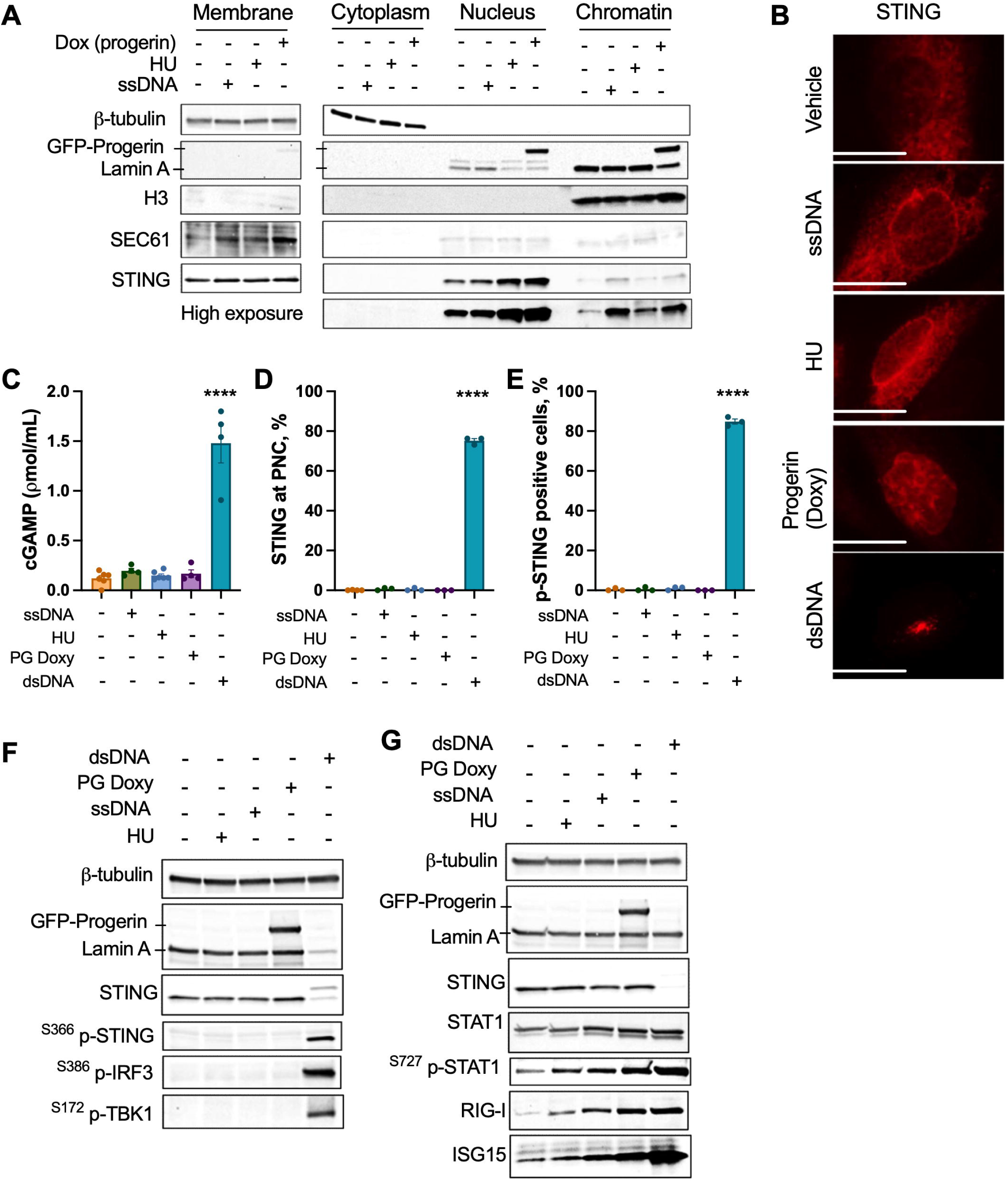
Replication stress induces nuclear accumulation of STING and activates an IFN response. (**A**) Western blot analysis of subcellular fractions from HDF treated for 6 days with doxycycline (Dox) to induce progerin expression, for 24 h with hydroxyurea (HU), or transfected with single-stranded DNA (ssDNA). Fraction markers include β-tubulin (cytoplasm), SEC61 (membrane), Lamin A (nucleus), and Histone H3 (chromatin). Note the increased presence of STING at nucleus and chromatin upon replication stress. Membranes imaged with prolonged exposure (high exposure) show a marked increase in signal intensity. (**B**) Immunofluorescence (IF) with STING antibody in HDF treated with vehicle, ssDNA, HU, Doxy (progerin), or dsDNA. (**C**) Quantification of cGAMP levels measured by ELISA in HDF treated as indicated. (**D**) IF with STING antibody and quantification of percentage of cells showing STING localization to the perinuclear compartment (PNC) upon different treatments. (**E**) IF with antibody recognizing phosphorylated STING on Ser366 and quantification of percentage of cells positive for ^S366^p-STING. (**F**) Immunoblot analysis of STING, GFP-progerin, and markers of activation of the canonical cGAS-STING pathway (^S366^p-STING, ^S386^p-IRF3, and ^S172^p-TBK1) following treatments. (**G**) Immunoblot analysis of STING pathway components and ISG proteins (STAT1, ^S727^p-STAT1, RIG-I, and ISG15) after indicated treatments.

We next asked whether RS-associated stimuli activate the canonical or non-canonical cGAS–STING pathways. Measurement of cGAMP levels showed that, unlike dsDNA, neither HU nor ssDNA induced a detectable increase in cGAMP production (**Figure 1C**). Similarly, STING phosphorylation (^S366^p-STING) and translocation to the PNC were observed only in response to dsDNA, not upon HU treatment, ssDNA transfection, or progerin expression (**Figure 1D–E**). Immunoblotting confirmed that only dsDNA activates downstream effectors such as TBK1 (^S172^p-TBK1) and IRF3 (^S386^p-IRF3), while replication stress stimuli (HU, ssDNA) failed to do so (**Figure 1F**). Despite this, HU and ssDNA still triggered an IFN response, like progerin-expressing cells, characterized by STAT1 phosphorylation and upregulation of ISGs such as ISG15 and RIG-I (**Figure 1G**), indicating that RS promotes IFN signaling even in the absence of canonical cGAS-STING pathway activation. Altogether, these data show that RS, whether induced by HU, mimicked by ssDNA transfection, or triggered by progerin, leads to nuclear accumulation of STING and chromatin association. However, it does not activate the canonical cGAS–STING pathway, suggesting a non-canonical mechanism by which STING contributes to inflammatory signaling in cells under chronic or endogenous replication stress.

### STING inhibition improves DNA damage, proliferation, and replication stress markers in progerin-expressing cells

The presence of STING in the nucleus raises questions about its role in genome maintenance. We confirmed that STING localizes to the nucleus in HDF, binding to lamin A, as shown by proximity ligation assay (PLA) with lamin A and STING antibodies (**Figure 2A**). Recent studies have revealed that nuclear STING is involved in gene transcription, DNA damage response regulation, and DNA replication ^11,31^. However, the mechanisms by which STING contributes to these processes remain poorly understood.

**Figure 2.**
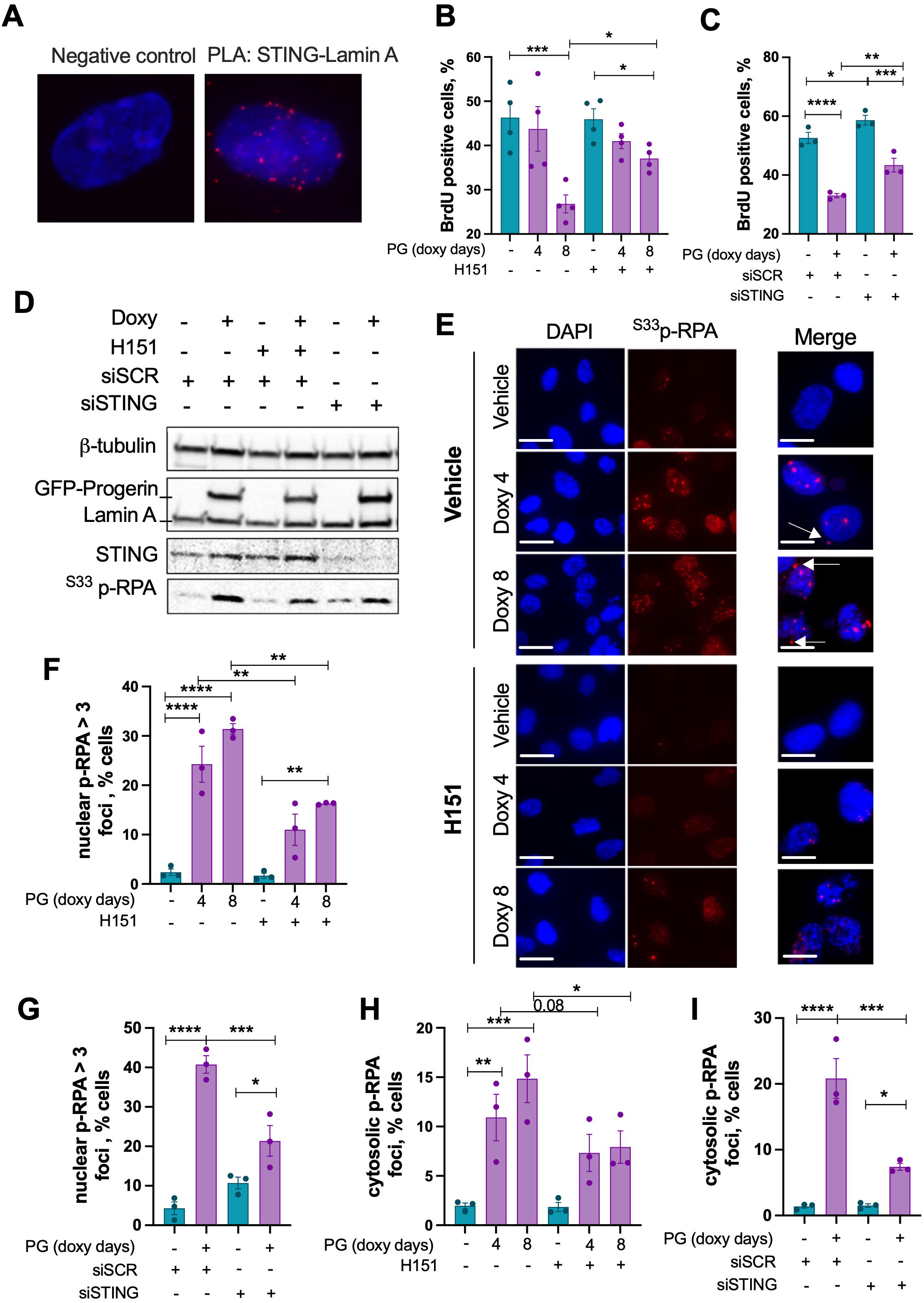
STING inhibition restores replication and suppresses replication stress markers in progerin-expressing cells. (**A**) Representative images of PLA on HDF shows STING-Lamin A interaction (foci) in control and progerin (PG)-expressing cells. DAPI is shown labeling genomic DNA. (**B**) Immunofluorescence analysis of BrdU incorporation in HDF treated with vehicle or doxycycline (Doxy) for 4 or 8 days to induce GFP-progerin, with or without STING inhibitor H151. (**C**) BrdU incorporation in HDF and progerin-expressing HDF transfected with STING siRNA (siSTING) or non-targeting control (siSCR). (**D**) Immunoblot analysis of STING, GFP-progerin, and phosphorylated RPA (^S33^p-RPA) in HDF control and progerin-expressing HDF in which STING is abrogated (H151 or siSTING). (**E**) IF images showing ^S33^p-RPA foci (red) in cells treated with Doxy ± H151 for 4 or 8 days; DAPI (blue) labels nuclei and white arrows indicate cytosolic foci. Scale bars, 20 µm (**F**) Quantification of cells with ≥3 nuclear ^S33^p-RPA foci per nucleus from (E). (**G**) Quantification of cells with nuclear ^S33^p-RPA foci in siSTING-or control siSCR-transfected cells treated with Doxy for 8 days. (**H**) Percentage of cells showing with ≥3 cytosolic ^S33^p-RPA foci from (E). (**I**) Cytosolic ^S33^p-RPA foci quantification in siSTING versus siSCR-transfected cells.

We tested a role for STING on genomic instability, particularly given our prior reports showing that progerin expression reduces RAD51 levels ^14^; a key factor in DSB repair by homologous recombination and in replication fork stability. As shown in **Figure S1A**, pharmacological inhibition of STING (H151) increases RAD51 protein levels, suggesting that STING might contribute to genomic instability in progeria. We then assessed markers of DNA damage. Prolonged progerin expression (4 and 8 days) led to progressive accumulation of γH2AX and 53BP1 foci, consistent with our previous findings ^32^ (**Figure S1B–C**). STING inhibition modestly reduced γH2AX- and 53BP1-positive cells, particularly at later timepoints, indicating a partial alleviation of DNA damage signaling. These findings support a role for STING in modulating the DNA damage response in progeria, though its inhibition is not sufficient to fully reverse genomic instability, suggesting that it acts in concert with other stress-related pathways.

STING has been implicated in DNA replication, mediating the protection of replication forks in some cases ^33^, and inducing replication fork instability in others ^31^. To test the functional consequences of STING nuclear accumulation in progerin-expressing cells, we tested whether its abrogation could improve proliferation and replication stress. Using HDF with inducible progerin expression, we either inhibited STING with H151 or depleted STING using specific siRNAs. Both approaches significantly enhanced cell proliferation, as shown by increased BrdU incorporation (**Figure 2B–C**). We then defined the extent of RS by monitoring RPA phosphorylation (^S33^p-RPA) by immunoblotting, the main marker of RS. We found that STING abrogation, through either H151 treatment or siRNA depletion, led to a marked reduction in ^S33^p-RPA (**Figure 2D** and **Figure S1A**). This effect was confirmed by immunofluorescence (**Figure 2E**). Note the presence of nuclear ^S33^p-RPA foci in HDF induced to express progerin for 4-8 days, and how these are reduced by STING inhibition with H151 (**Figure 2F-G**). Additionally, progerin-expressing HDF frequently exhibited aberrant cytoplasmic localization of ^S33^p-RPA, which was significantly reduced upon STING targeting **(Figure 2H–I**). These results demonstrate that STING contributes to replication stress, impaired proliferation, and self-DNA buildup in the cytosol (^S33^p-RPA-bound ssDNA) in progeria cells, and that its inhibition, whether pharmacologically or genetically, can partially restore cell cycle progression and mitigate RS markers.

### STING causes replication fork slowing/stalling and nascent DNA degradation in progerin-expressing cells

Recently, a function of STING promoting nascent DNA degradation (NDD) in the context of cancer was reported ^31^. To determine whether STING directly associates with nascent DNA in HDF and HDF-progerin, we performed SIRF (*in situ* protein interaction with nascent DNA at replication forks) ^34^. We find PLA signals in HDF and HDF-progerin cells (**Figure 3A**), demonstrating the binding of STING protein to nascent DNA (EdU-biotin labeled) at the replication fork. The role that STING plays in replication remains unknown.

**Figure 3.**
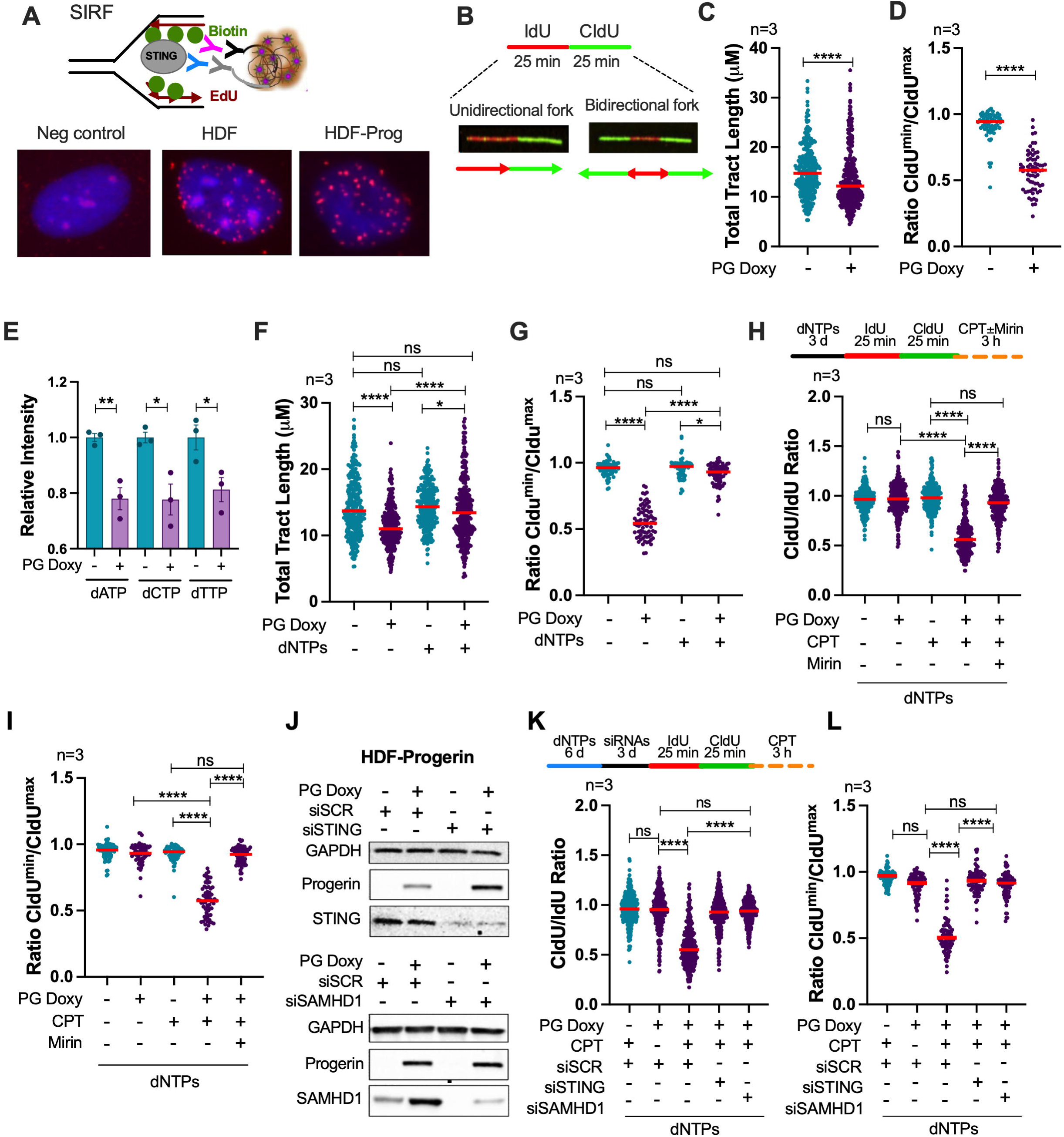
Prolonged progerin expression leads to replication fork slowing and degradation. (**A**) SIRF performed in HDF control and progerin-expressing HDF. Note the localization of STING in close proximity to EdU-biotin labeled nascent DNA. (**B**) Schematic representation of DNA fiber assay labeling strategy showing unidirectional and bidirectional fork progression. Cells were sequentially labeled with IdU (red) and CldU (green) for 25 minutes each. Bidirectional forks appear as green-red-green tracks extending from a common origin. (**C**) Quantification of total tract length (IdU + CldU) indicating fork speed, in control and progerin-expressing (Dox 6 days) HDF. (**D**) Quantification of fork symmetry expressed as the ratio of the shorter CldU track to the longer CldU track in each bidirectional fork (CldU^min^/CldU^max^). (**E**) Quantification of dATP, dCTP, and dTTP in control and progerin-expressing HDF by mass spectrometry. (**F**) DNA fiber assay quantifying total tract length in progerin-expressing cells supplemented or not with dNTPs (1X) for 3 days prior labeling. (**G**) Fork symmetry analysis from (F) under the same conditions. (**H**) Replication fork instability (RFI) analysis in control and progerin-expressing cells that are supplemented with dNTPs. We performed labeling of nascent DNA with IdU + CldU, followed by treatment with camptothecin (CPT, 3 hours) to stall the forks. CldU/IdU ratios were used to measure RFI, and Mirin was added to determine whether RFI in progerin-expressing cells was due to fork degradation. (**I**) Fork symmetry measurements under the same treatment conditions as in (H). For all quantifications, each point represents a single replication fork; n = 3 biologically independent experiments. (**J**) Immunoblots showing STING knockdown (top) and SAMHD1 (bottom) in HDF control and progerin-expressing HDF. (**K**) DNA fiber analysis in control and progerin-expressing HDF that were supplemented with dNTPs on day 0, transfected with siRNAs (siSTING or siSAMHD1) on day 3, and DNA fibers performed on day 6 (with or without addition of CPT). (**L**) Fork symmetry from the same samples as in (K).

To investigate the functional link between STING and progerin in DNA replication, we analyzed replication fork dynamics using DNA fiber assays after 6 days of progerin induction; a time point characterized by STING upregulation and IFN response activation ^16^. We monitored replication fork dynamics by pulse labeling cells with different thymidine analogs (IdU and CldU for 25 minutes each), performed DNA fiber assays ^35^ and quantified the total tract length of unidirectional forks and the sister CldU tracts length in the bidirectional fibers (**Figure 3B**). The total tract length was measured as an indicator of fork speed. Our data show that, after 6 days of progerin expression and compared to control cells, replication speed is significantly slowed, as indicated by the reduced total tract length of fibers (**Figure 3C**). Additionally, we observed increased asymmetry in the sister CldU tracts of bidirectional forks (displayed as a ratio of the shorter CldU tract to the longer CldU tract of the same sister fork; CldU^min^:CldU^max^) (**Figure 3D**) in progerin-expressing cells. This fork asymmetry indicates an increase in fork stalling in progerin-expressing cells.

One mechanism that could explain fork slowing and stalling is the impaired nucleotide metabolism reported previously in HGPS patients-derived cells ^36^. Metabolomics performed in HDF-control and HDF-progerin confirmed that expression of progerin reduces significantly the levels of dNTPs (**Figure 3E**). To test whether this deficiency underlies RS, we added exogenous dNTPs after the induction of progerin, and 3 days before performing DNA fiber assays. Supplementation of dNTPs restored average fork speed and significantly corrected fork asymmetry (**Figure 3F–G**). These data indicate that insufficient nucleotide pools underlie the RS phenotype in progerin-expressing cells, and that STING impacts dNTP metabolism in this context.

We next tested whether dNTP-replenished progeria cells could protect stalled replication forks from degradation; as replication fork instability (RFI) upon stalling is a phenotype characteristic of progeria cells ^13,28,29^. Fork stalling was induced after IdU+CldU labeling using camptothecin (CPT), which inhibits topoisomerase I without altering nucleotide levels. While dNTP supplementation restored fork progression under unperturbed conditions, it failed to prevent nascent DNA degradation (NDD) following CPT-induced stalling, as evidenced by a reduced CldU/IdU ratio and increased sister fork asymmetry (**Figure 3H–I**). Treatment with the MRE11 inhibitor, Mirin, rescued the CldU/IdU ratio and restored symmetry (**Figure 3H–I**), confirming that fork degradation was mediated by this nuclease. These findings suggest that progerin-expressing cells suffer from both fork slowing due to dNTP depletion and a failure to protect stalled forks from MRE11-mediated degradation.

Given that STING inhibition reduces markers of RS (**Figure 2**) and that STING induces NDD in cancer cells ^31^, we determined whether STING contributes to RS phenotypes of progerin-expressing cells. In addition, we interrogated SAMHD1, a dNTPase and ISG that acts as a STING downstream effector. SAMHD1 plays a dual role in replication, acting as the main dNTPase that controls dNTPs levels during the cell cycle ^26^, while mediating the recruitment of MRE11 to stalled forks, facilitating NDD ^27^. Given that these two RS phenotypes are observed in progerin-expressing cells, that SAMHD1 is an ISG regulated by STING, and that it is upregulated in HGPS patients cells ^13^, we tested whether it mediates STING’s contribution to RS. First, we monitored the involvement of STING and SAMHD1 in NDD of stalled forks in progerin-expressing cells, by depleting the proteins (siRNAs) and performing DNA fiber assays (**Figure 3J**). To overcome dNTPs deficiency, and thus fork slowing and stalling, we performed these experiments in the presence of supplemented dNTPs since day 0. We induced the expression of progerin at day 0, depleted STING or SAMHD1 (siRNAs) at day 3 and performed labeling with IdU/CldU at day 6. After IdU/CldU labeling, RF stalling was induced by CPT treatment for 3 hours. Efficient STING and SAMHD1 knockdown were confirmed by immunoblotting (**Figure 3J**). To monitor NDD, we calculated the CldU/IdU ratio in the different conditions (**Figure 3K**). We find that progerin-expressing cells with supplemented dNTPs do not show a RFI phenotype, similarly to control HDF treated with CPT. However, when progerin-expressing HDF are treated with CPT, a clear RFI phenotype is observed. Importantly, depletion of STING or SAMHD1 prevented NDD in progerin-expressing cells, as evidenced by the ratio CldU/IdU=1 (**Figure 3K**), and rescued fork symmetry (**Figure 3L**). These data demonstrate that STING and SAMHD1 cause MRE11-mediated NDD of stalled forks in progerin-expressing cells, unveiling a toxic STING-SAMHD1 axis that drives RS and genomic instability in progeria.

### Expression of STING in tumor cells causes replication stress in a SAMHD1-dependent manner

To determine whether the involvement of STING in RS is conserved in different cellular contexts, we used U2OS cells, an osteosarcoma cell line that naturally lacks STING expression. We generated U2OS cell lines that constitutively express progerin, mCherry-STING, or both, and compared them to U2OS cells transfected with empty vector as control (U2OS-EV). This strategy allowed us to define the impact of STING alone in replication, as well as the role of STING in progerin-induced RS. Of note, we isolated different clones of U2OS with different levels of mCherry-STING expression and chose a clone with lower expression (lower mCherry intensity) to perform the studies. First, we asked whether STING is required for IFN signaling in this context. In U2OS cells, progerin expression alone failed to induce expression of interferon-stimulated genes (ISGs) such as ISG15 and RIG-I (**Figure 4A**). Overexpression of STING (mCherry-STING) did not induce ISGs either but increased the marker of replication stress ^S33^p-RPA. However, overexpression of STING and progerin induced the expression of these ISGs, along with an increase in ^S33^p-RPA (**Figure 4A**). To assess the role of STING in replication fork dynamics more directly, we performed DNA fiber assays. Expression of progerin alone had a minor impact on replication fork speed (**Figure 4B**) and modestly increased fork asymmetry compared to empty vector controls (**Figure 4C**). In contrast, STING overexpression, either in control U2OS cells or in combination with progerin, significantly reduced total tract length (**Figure 4B**) and increased fork asymmetry (**Figure 4C**), mimicking the RS phenotype seen in STING-expressing HDF-progerin cells. Note that the increased fork asymmetry is more pronounced when progerin and STING are co-expressed. These data demonstrate that increased expression of STING causes replication fork slowing/stalling also in tumor cells, and that the combination of progerin and STING expression exacerbates RS and leads to activation of an IFN response.

**Figure 4.**
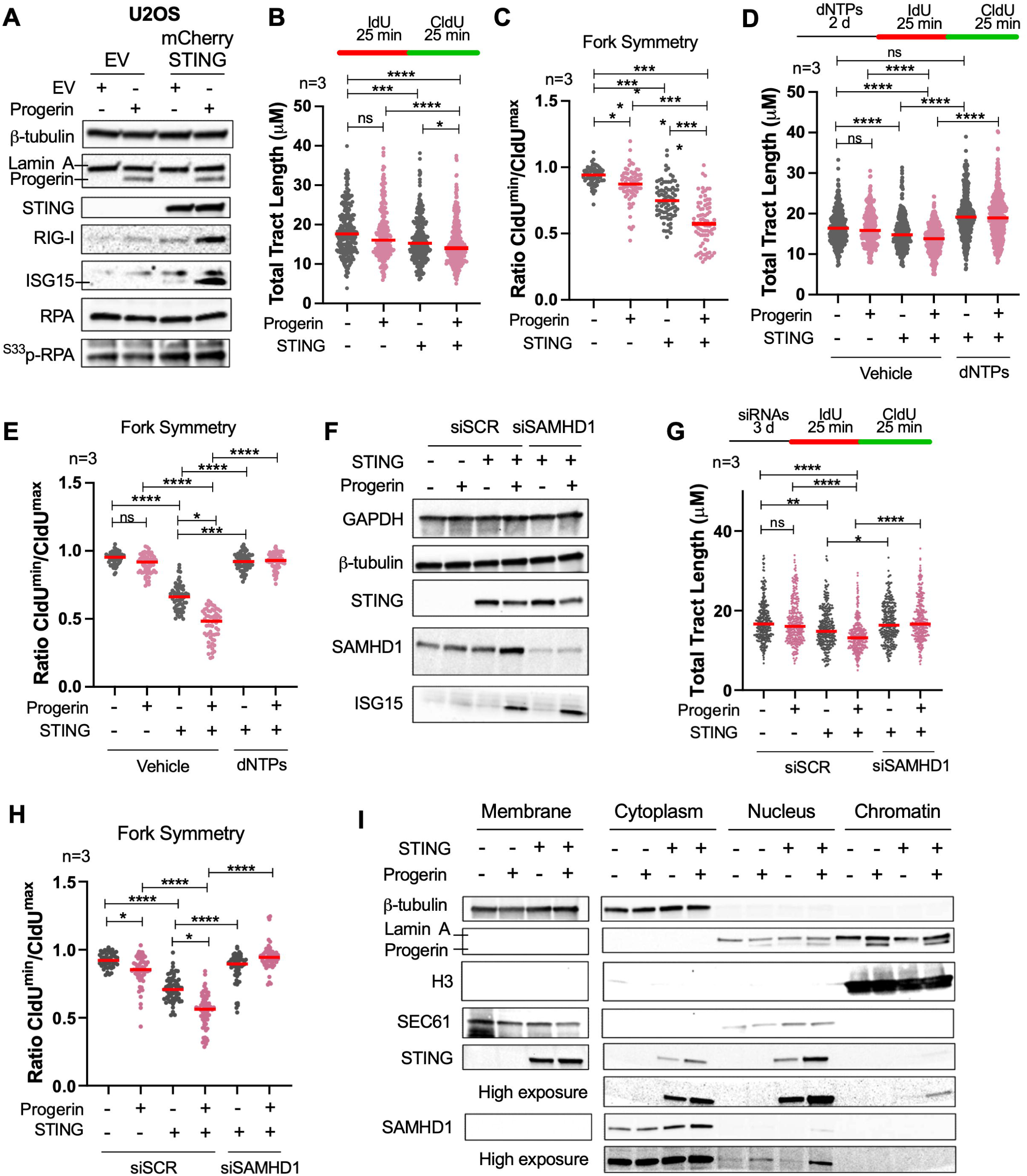
STING promotes replication fork instability in U2OS cells via SAMHD1. (**A**) U2OS tumor cells were generated that express progerin, STING, or both, using empty vectors (EV) as controls. Immunoblots performed to assess replication stress marker ^33^p-RPA, and activation of an IFN response (ISG15, RIG-I). (**B**) DNA fiber assays to monitor total tract length in U2OS +/- progerin and +/- STING. (**C**) Fork symmetry (ratio of the sisters CldU tracts from bidirectional forks) in same conditions as in (B). (**D**) DNA fiber analysis to monitor total tract length in U2OS +/- progerin and +/- STING with or without dNTP supplementation. (**E**) Fork symmetry in the same samples as in (D). (**F**) Immunoblot showing STING expression and SAMHD1 knockdown in control and progerin-expressing U2OS cells. (**G**) DNA fiber analysis to monitor total tract length in U2OS +/- progerin and +/- STING with or without SAMHD1 depletion. (**H**) Fork symmetry in the same samples as in (G). (**I**) Subcellular fractionation of U2OS +/- progerin and +/- STING, probed for STING and SAMHD1. H3 and SEC61 were used as nuclear and ER markers, respectively. For all quantifications, each point represents a single replication fork; n = 3 biologically independent experiments.

Next, we tested whether the replication defects observed in U2OS STING-expressing cells could be directly linked to dNTP availability. Remarkably, upon dNTP supplementation, replication fork speed was restored in cells expressing STING and in cells expressing both STING+progerin, as shown by increased total tract lengths (**Figure 4D**). Furthermore, fork stalling was also significantly reduced in all contexts, as shown by improved fork symmetry (**Figure 4E**). These results indicate that elevated STING levels impair replication dynamics through a mechanism that can be rescued by restoring nucleotide pools.

To determine whether SAMHD1 is part of the mechanism leading to the replication stress induced by STING in U2OS cells, we performed DNA fiber assays upon depletion of SAMHD1 by siRNA. As shown in **Figure 4F**, we achieved a marked reduction of SAMHD1 in U2OS cells. DNA fiber assays show that the reduced total tract length and the reduced asymmetry observed in STING- and STING+progerin-expressing cells was rescued by SAMHD1 depletion (**Figure 4G-H**). These data confirm that SAMHD1 is necessary for the RS phenotype induced by STING in U2OS cells, mirroring its function in progerin-expressing fibroblasts.

Moreover, we performed subcellular fractionation to determine how STING expression in U2OS cells impacts the levels and localization of SAMHD1. We find that both STING and SAMHD1 are present in cytoplasm and nuclear fractions, especially upon progerin expression (**Figure 4I**). SAMHD1 is localized primarily in the cytoplasm in all contexts, however an increased presence in the nucleus is observed upon expression of both STING and progerin. Together, these results support a model in which STING promotes replication fork instability through SAMHD1 by impairing fork progression and promoting NDD.

## Discussion

STING is an adaptor protein involved in cytosolic DNA sensing, mediating the canonical cGAS–STING pathway through its translocation to the perinuclear compartment and subsequent activation of TBK1 and IRF3 ^37,38^. However, emerging studies, including our own, reveal that STING participates in non-canonical signaling pathways ^19,39^, and is present in the nucleus, where its functions remain understudied. Here, we show that inducing RS via dNTPs depletion (HU), or mimicking RS with ssDNA transfection, trigger STING nuclear accumulation and chromatin binding, mirroring progerin-expressing cells. This shift in STING from cytosolic signaling to chromatin binding implies that under endogenous genomic stress conditions, STING engages in non-canonical nuclear pathways. These results are aligned with recent studies proposing nuclear functions for STING in transcriptional regulation, chromatin remodeling, and nascent DNA metabolism at stalled forks ^23,31,40^ in some contexts, although the precise mechanisms remain to be elucidated.

Our study demonstrates that activation of the non-canonical cGAS-STING pathway in progerin-expressing cells (∼6 days after doxycycline-induced progerin expression) hinders DNA replication via mechanisms that involve STING and its downstream effector SAMHD1. We propose a model whereby progerin expression rapidly causes genomic instability and nuclear fragility ^13,14,29^, leading to cytosolic DNA buildup and activation of the non-canonical nuclear STING pathway, featuring increased expression of ISGs. Several ISGs are known to influence replication, including ISG15 ^41,42^, APOBEC3 proteins ^43,44^ and SAMHD1 ^26,27^. Of these, SAMHD1 emerges as the key contributor to dNTPs depletion, which causes RF slowing and stalling. In this manner, STING indirectly regulates replication fork dynamics, adding a novel layer of complexity to STING’s function beyond immunity. Another function of SAMHD1 is the facilitation of MRE11 recruitment and activity at stalled forks, which leads to nascent DNA resection and processing, necessary for proper fork restart ^27^. This function of SAMHD1 is protective, limiting the amount of ssDNA fragments released into the cytosol during RS, which prevents activation of cGAS-STING and the IFN response ^27^. Accordingly, depletion of SAMHD1 leads to activation of the IFN response in a variety of cell lines, in a cGAS- and STING-dependent manner ^27^. Also, mutations in SAMHD1 are associated with an interferonopathy known as Aicardi-Goutieres syndrome (AGS). However, we find that this function of SAMHD1 has detrimental effects for replication in the context of progerin expression and STING upregulation and nuclear/chromatin accumulation. The increased levels of SAMHD1 in progerin-expressing cells might lead to increased recruitment of MRE11 to the stalled forks and their excessive processing and degradation. Thus, the STING-SAMHD1 axis constitutes a pathological feedforward loop, further exacerbating hallmarks of aging such as genomic instability and inflammation. Targeting this pathway either through STING or SAMHD1 inhibition may break this cycle, offering a promising therapeutic strategy to alleviate both replication defects and chronic inflammation.

This model of a toxic STING-SAMHD1 axis causing RS is recapitulated in U2OS tumor cells, which normally lack STING expression. Unexpectedly, expression of progerin alone in U2OS cells only causes a modest RS phenotype, which is exacerbated upon STING expression, reinforcing the idea that progerin-induced RS phenotypes require STING. Importantly, we show that STING expression alone has a toxic effect in replication in U2OS cells, which is also mediated by SAMHD1. These findings support recent work showing that STING regulates the fate of stalled forks through RPA loading, S-phase checkpoint regulation, and promotion of NDD ^31^. STING also fractionated to nuclear fractions such as chromatin and nuclear envelope ^31^. Thus, these two studies support the idea that STING is a determinant of replication fork stability, with important implications for chemotherapeutic strategies that stall replication of tumor cells. The presence or absence of STING in this scenario might determine the response to replication poisons and might also be behind the loss of STING in a wide variety of tumor cells ^45^. Loss of STING in tumor cells could provide a survival and proliferative advantage, reducing NDD and genomic instability, in addition to its well-known effect enabling tumor cells to evade the immunosurveillance system. Future work needs to determine whether STING binding to nascent DNA, as shown here, plays a direct role in replication. We envision that STING might act as a platform for the recruitment of SAMHD1-MRE11 complex or other replication fork remodeling and protective factors. Moreover, understanding how other age-associated and cancer-promoting pathways intersect with STING-SAMHD1-mediated RS will be crucial for designing strategies to ameliorate genomic instability and inflammation in disease contexts.

In conclusion, our study reveals a multi-tiered mechanism by which STING integrates metabolic and enzymatic pathways to orchestrate the cellular response to replication stress in progeria and tumor cells. In these contexts, STING acts as an active participant in genome destabilization, linking metabolic imbalance and immune activation in a cohesive pathological program. These findings suggest that STING inhibition could serve as a dual-purpose therapy, restoring replication fidelity and dampening inflammation in progeria and potentially other aging-related diseases marked by chronic RS and innate immune activation. In tumor cells however, restoring STING function could be used as a strategy to induce RS and genomic instability, while restoring anti-tumor immunity. Defining mechanistically the different functions of STING will allow us to design better strategies to manipulate STING with therapeutic benefits for aging and cancer.

## Material and Methods

### Cell culture and cell lines

Human dermal fibroblasts containing doxycycline-inducible GFP-progerin construct were cultured at 37°C in DMEM supplemented with 10% FBS and 1% antibiotics, antimycotics and treated for 4,6 and 8 days with vehicle (DMSO) or 1µg/mL doxycycline. U2OS cells containing progerin (PG) or empty vector (EV) were also cultured using the same conditions.

### cGAMP assessment

For cGAMP measurement, cells transfected with 1μg/ml of poly (dA:dT) for 4 hr were lysed in RIPA buffer followed by centrifugation at 14000 x g at 4°C for 15 min. The supernatant was used for cGAMP ELISA assay according to manufacturer’s instructions (2’3’- Cyclic GAMP ELISA Kit, ThemoFisher). cGAMP concentration is shown for 100 μg of protein.

### BrdU incorporation assay

For 5-bromo-2’-deoxyuridine (BrdU) incorporation assay as a measurement of cell proliferation, cells were seeded onto sterile glass coverslips in 24-well plates and allowed to adhere overnight. The following day, BrdU was added to the culture medium at a final concentration of 10 µM and incubated for 5 hours at 37°C under standard culture conditions. Following BrdU labeling, cells were subjected to a pre-extraction step to remove soluble proteins and enhance nuclear signal detection. Cells were incubated with pre-extraction buffer (20mM HEPES, 50mM NaCl, 3mM MgCl2, 0.5% Triton-X-100 in phosphate-buffered saline [PBS]) for 10 minutes on ice. Immediately afterward, cells were fixed with 4% paraformaldehyde (PFA) in PBS for 20 minutes at room temperature. After three 5-min washes with PBS, DNA was denatured by incubating cells with 2N hydrochloric acid (HCl) for 20 minutes at room temperature, followed by neutralization with 0.1 M sodium borate buffer (pH 8.5) for 10 minutes. This wash was repeated three times. After three washes in PBS, non-specific binding was blocked by incubation with 2% bovine serum albumin (BSA) in PBS for 30 minutes at room temperature, and the immunofluorescence using anti-BrdU antibody was performed as previously described.

### Analysis of dNTP levels

The levels of dNTP were analyzed in HDF^tetON^-GFP-progerin cells 6 days after starting the treatment with vehicle or Doxy (1 µg/ml). 24 hours prior the collection of cells the medium was changed to a fresh growth medium without vehicle or Doxy. On day of collection the cells were washed twice with PBS followed by two washes with LC-MS grade water and then quenched with ice-cold LC-MS grade methanol. The cells were scraped, collected into microcentrifuge tubes, and stored at -80 °C until further analysis. The samples were submitted to the Center for Mass Spectrometry and Metabolic Tracing at Washington University for dNTP analysis. The metabolomics sample preparation and LC/MS analysis were performed as reported previously^46^.

### DNA fibers assay

The specific labeling scheme and treatments for each experiment are shown in the legends or schematics in the figures. When mentioned, cells were supplemented with exogenous dNTP (Embryomax^®^ Nucleosides 100×□- Merck, ES-008-D) that were added to cell culture media at a final concentration of 1×□for 3 or 6 days as indicated in the figures^36^. Cells were plated one day before performing the labeling. Asynchronous cells were labeled with Thy analogs: 20 μM IdU followed by 200 μM CldU. Cells were collected by trypsinization, washed, and resuspended in 70μl of PBS. Then, 2 μl of cell suspension was dropped on positively charged slide (Denville Ultraclear), and cell lysis was performed in situ by adding 8 μl lysis buffer (200 mM Tris–HCl [pH 7.5]; 50 mM EDTA; and 0.5% SDS). Stretching of high-molecular weight DNA was achieved by tilting the slides at 15 to 45°. The resulting DNA spreads were air dried and fixed for 5 min in 3:1 methanol:acetic acid and refrigerated at 4 °C. For immunostaining, stretched DNA fibers were denatured with 2.5 N HCl for 60 min, washed 3× for 5 min in PBS, then blocked with 5% bovine serum albumin (BSA) in PBS for 30 min at 37 °C. Mouse anti-IdU/BrdU (BD Biosciences; 347580) (1:20), and goat antimouse IgG1 Alexa 547 (Invitrogen; A21123) (1:100) and Rat anti-CldU/BrdU (Abcam; ab6326) (1:75), chicken antirat Alexa 488 (Invitrogen; A21470) (1:100) antibodies were used to reveal IdU-labeled and CldU-labeled tracts, respectively. Labeled tracts were visualized under a Leica DM5000B microscope using 63× oil objective lens with a Lecia DFC350FX digital camera and Lecia application Suite. The tract lengths were measured using ImageJ (the National Institutes of Health). A total of 100 to 200 fibers were analyzed for each condition in each experiment. All analyses were performed blinding the samples. Statistical analysis of the tract length was performed using an unpaired two-tailed t-test; Mann-Whitney or *Kruskall-Wallis* test followed by Dunn’s multiple comparisons on GraphPad Prism (GraphPad Software, Inc).

### Immunoblotting

Immunoblotting was carried out by lysing cells in RIPA buffer (Thermo Scientific- 89901), for 30min rotating at 4°C. Lysates were centrifuged at 14,000rpm for 12min at 4°C and pellet was removed from the sample. Following centrifugation, 50-60 µg of total protein in the supernatants were mixed with Laemmli loading buffer and incubated at 95°C for 5 min prior electrophoresis. Equal protein amounts were separated by SDS-PAGE on a 4-15% Criterion TGX Gel (Bio-Rad) followed by transferring to a nitrocellulose membrane using the Trans-Blot Turbo system (Bio-Rad). Membranes were blocked using 5% BSA in PBS + 0.1% Tween-20 for 1 h at room temperature and then incubated overnight at 4°C with the appropriate primary antibody diluted in blocking solution. Primary antibodies used were β-tubulin (1:5000- Origene-AP31823PU-N), GAPDH (1:1000- Cell Signaling-2118), Lamina A (1:3000- Abcam-1791), Progerin (1:1000 Santa Cruz-81511), ISG15 (1:1000 Santa Cruz-166755), STING (1:1000-Cell Signaling-13647), ^S366^p-STING (1:1000- Cell Signaling- 50907), RIG-I (1:1000-Cell Signaling-3743S), S33-p-RPA (1:1000 Bethyl- PLA0070), SAMHD1 (1:1000- Cell Singaling-49158), λH2AX (1:1000- Cell Signaling- 2577). Membranes were washed three times using PBS + 0.1% Tween-20 after both primary and secondary antibody incubations. Membranes were developed using Pierce ECL Western Blotting Substrate (Thermo Scientific-32132). Western blot images were acquired using SYNGENE PXi instrument.

### Immunofluorescence

Immunofluorescence was performed to analyze STING localization and activation (STING 1:500-Cell Signaling- 13647, ^S366^pSTING 1:1200- Cell Signaling-50907), BrdU quantification (1:200- Santa Cruz -sc20045), levels of ^S33^p-RPA (1:1000- Bethyl -PLA0070), γH2AX (1:200 Cell Signalling-2577) and 53BP1 (1:1000- Santa Cruz- sc517281). Cells were plated onto coverslips and allowed to fully attach and were submitted to different treatments according to text or each figure legend. After treatment, cells were washed and placed on ice for 10 minutes in an extraction buffer containing 25 mM Hepes (pH 7.4), 50 mM NaCl, 1 mM EDTA, 3 mM MgCl_₂_, 300 mM sucrose, and 0.5% Triton X-100 in double-distilled water. Cells were then washed three times with 1× PBS, fixed in 4% formaldehyde for 10 minutes (fixation with methanol for 10 minutes was used for STING antibody), permeabilized with 0.5% Triton X-100 for 30 minutes at room temperature, and blocked for 1 hour at 37°C in 1% BSA/PBS. Primary antibody incubation was carried out overnight at 4°C, followed by secondary antibody incubation for 1 hour at 37°C, both in a humidified chamber. After PBS washes, cells were counterstained with DAPI and mounted in Vectashield. For detection of STING and phospho-STING (p-STING) cells were fixed in ice-cold methanol (−20 °C) and then permeabilization, blocking and incubation with primary and secondary antibodies were performed as described above. Imaging was performed using a Leica DM5000B microscope with 40× or 63× oil-immersion objectives, and photos were taken with a Leica DFC350FX camera using the Leica Application Suite. The primary antibody used was phospho-RPA at serine 33 (Bethyl, A300-246A).

### Proximity ligation assay

HDF^TetON^-GFP-progerin cells were synchronized with aphidicolin (1□µM) overnight. The following day, the medium was replaced to remove aphidicolin and allow cells to resume DNA replication. For SIRF-PLA, cells were then pulsed with EdU (10 µM) for 1h to label nascent DNA. After EdU incorporation, cells were washed with PBS (1×) and pre-extracted on ice for 5 min using pre-extraction buffer (20 mM HEPES, 50 mM NaCl, 3 mM MgCl_₂_, 0.5% Triton X-100). Cells were subsequently fixed with 4% formaldehyde at room temperature and washed three times with PBS (1×). The Click-iT reaction was performed in PBS containing 10 µM biotin-azide, 2 mM CuSO_₄_, and 10 mM sodium ascorbate for 30 min at room temperature in the dark. Following the click reaction, cells were blocked with 10% goat serum in PBS for 1 h at 37 °C, then incubated overnight at 4 °C with primary antibodies against STING (Invitrogen; MA5-26030) (1:100) and anti-biotin (Cell signaling; 5597S) (1:200). After washing, proximity ligation was carried out according to the manufacturer’s instructions (Duolink® In Situ Red Kit Mouse/Rabbit (Sigma-Aldrich, Cat # DUO92008). For Lamin A–STING PLA, cells were processed as described above, omitting the EdU labeling step. Cells were incubated with anti-STING and anti-Lamin A (Abcam; ab26300) (1:500). Images were acquired using a Leica DM5000B microscope using 63× oil objective lens with a Lecia DFC350FX digital camera and Leica application Suite. STING-EDU PLA foci were quantified using CellProfiler 4.2.5 software and ImageJ 1.54g software was used to quantify STING-LMNA PLA foci

### RNA interference

Transient transfection of siRNAs was performed using Lipofectamine™ 3000 Transfection Reagent (Invitrogen- L3000015). SMARTpool siRNA Dharmacon was used to deplete STING (L-024333-00-0005; 50nM; 72 h), SAMHD1 (L-013950-01-0005; 50 nM, 72h). In each experiment, non-targeting siRNA (Dharmacon- ID:D-001810-10-05) was used as transfection control.

### Subcellular fractionation

HDF cells were exposed to 1 mM HU or transfected with 2 µg/ml ssDNA for 24 h prior to subcellular fractionation. ssDNA transfection was performed using Lipofectamine™ 3000 Transfection Reagent (Invitrogen- L3000015). HDF^TetON^-GFP-progerin were treated for 6 days with vehicle (DMSO) or 1µg/mL doxycycline. Subcellular fractionation was performed using “Subcellular protein fractionation kit for cultured cells” from Fisher Thermo Scientific according to the manufacturer’s instructions.

### Lentiviral transduction

Virus production and transduction were carried out in three stages as described in^35,47,48^. First, 293T producer cells were co-transfected with (i) the VSV-G envelope plasmid pCMV-VSV-G, (ii) a packaging helper (pUMVC3 for retrovirus or pHR’8.2ΔR for lentivirus) and (iii) one of three transfer vectors: pLVBsd-CMV-mCherry-hSTING, pMXIH-G608G (progerin), or the matching empty control pMXIH-EV. Forty-eight hours later, the conditioned medium containing viral particles was collected and in the second stage, U2OS target cells were transduced with the supernatant for two consecutive days to maximize infection efficiency. Finally, transduced cultures were allowed a 24–48 h recovery period before antibiotic selection, hygromycin or neomycin, depending on the vector backbone, was applied to enrich for successfully infected cells.

### Quantification and statistical analysis

Statistical analysis was performed using Prism 8 (GraphPad Software). Details of the individual statistical tests are indicated in the figure legends and results. All experiments were repeated at least three times unless otherwise noted. Statistical differences in DNA fiber tract lengths were determined by Kruskal–Wallis followed by Dunn’s multiple comparisons test. Statistical analysis for immunofluorescence assays was determined by two-way ANOVA followed by Bonferroni test.

## Author Contributions

Barbara Teodoro-Castro and Rafael Cancado de Faria contributed equally, performing most of the experiments, and writing the first draft of the manuscript. Elena V. Shashkova and Atika Malique performed some experiments. Madison Adolph provided expertise guidance on PLA, STING binding to nascent DNA by SIRF, and overall discussion of results and conclusions. Lilian N. D. Silva performed some experiments and supervised the work and the preparation of the manuscript for submission. Susana Gonzalo supervised the work.

## Acknowledgments

This work was supported by the National Institutes of Health [RAG082759A, RAG076145A, and R01AG058714 to S.G.]; the Glenn Foundation for Medical Research Postdoctoral Fellowship in Aging Research [PD24164 to L.N.D.S.]; and the Edward A. Doisy Department of Biochemistry and Molecular Biology Scholarship to R.C.F., B.T.C., and A.M.

## References

1 Wu, J. et al. Cyclic GMP-AMP is an endogenous second messenger in innate immune signaling by cytosolic DNA. Science 339, 826–830, doi:10.1126/science.1229963 (2013).

2 Ivanov, A. et al. Lysosome-mediated processing of chromatin in senescence. J Cell Biol 202, 129–143, doi:10.1083/jcb.201212110 (2013).

3 Dou, Z. et al. Cytoplasmic chromatin triggers inflammation in senescence and cancer. Nature 550, 402–406, doi:10.1038/nature24050 (2017).

4 Harding, S. M. et al. Mitotic progression following DNA damage enables pattern recognition within micronuclei. Nature 548, 466–470, doi:10.1038/nature23470 (2017).

5 Mackenzie, K. J. et al. cGAS surveillance of micronuclei links genome instability to innate immunity. Nature 548, 461–465, doi:10.1038/nature23449 (2017).

6 Gluck, S. et al. Innate immune sensing of cytosolic chromatin fragments through cGAS promotes senescence. Nat Cell Biol 19, 1061–1070, doi:10.1038/ncb3586 (2017).

7 Takahashi, A. et al. Downregulation of cytoplasmic DNases is implicated in cytoplasmic DNA accumulation and SASP in senescent cells. Nat Commun 9, 1249, doi:10.1038/s41467-018-03555-8 (2018).

8 Ahn, J. et al. Inflammation-driven carcinogenesis is mediated through STING. Nat Commun 5, 5166, doi:10.1038/ncomms6166 (2014).

9 Yang, H., Wang, H., Ren, J., Chen, Q. & Chen, Z. J. cGAS is essential for cellular senescence. Proc Natl Acad Sci U S A 114, E4612–E4620, doi:10.1073/pnas.1705499114 (2017).

10 Zheng, W. et al. The Role of cGAS-STING in Age-Related Diseases from Mechanisms to Therapies. Aging Dis 14, 1145–1165, doi:10.14336/AD.2023.0117 (2023).

11 Zhang, R. et al. Nuclear localization of STING1 competes with canonical signaling to activate AHR for commensal and intestinal homeostasis. Immunity 56, 2736–2754 e2738, doi:10.1016/j.immuni.2023.11.001 (2023).

12 Gulen, M. F. et al. cGAS-STING drives ageing-related inflammation and neurodegeneration. Nature 620, 374–380, doi:10.1038/s41586-023-06373-1 (2023).

13 Kreienkamp, R. et al. A Cell-Intrinsic Interferon-like Response Links Replication Stress to Cellular Aging Caused by Progerin. Cell Rep 22, 2006–2015, doi:10.1016/j.celrep.2018.01.090 (2018).

14 Coll-Bonfill, N., Cancado de Faria, R., Bhoopatiraju, S. & Gonzalo, S. Calcitriol Prevents RAD51 Loss and cGAS-STING-IFN Response Triggered by Progerin. Proteomics 20, e1800406, doi:10.1002/pmic.201800406 (2020).

15 de Faria, R. C. & Gonzalo, S. Sterile inflammation in laminopathies. Eur J Cell Biol 104, 151512, doi:10.1016/j.ejcb.2025.151512 (2025).

16 Cancado de Faria, R., et al. STAT1 Drives Interferon-Like Response and Aging Hallmarks in Progeria. Aging Biology, 1–15, doi::10.59368/agingbio.20230009 (2023).

17 Ishikawa, H., Ma, Z. & Barber, G. N. STING regulates intracellular DNA-mediated, type I interferon-dependent innate immunity. Nature 461, 788–792, doi:10.1038/nature08476 (2009).

18 Ablasser, A. et al. cGAS produces a 2’-5’-linked cyclic dinucleotide second messenger that activates STING. Nature 498, 380–384, doi:10.1038/nature12306 (2013).

19 Cancado de Faria, R., et al. A noncanonical cGAS-STING pathway drives cellular and organismal aging. Proc Natl Acad Sci U S A 122, e2424666122, doi:10.1073/pnas.2424666122 (2025).

20 Guey, B. & Ablasser, A. Emerging dimensions of cellular cGAS-STING signaling. Curr Opin Immunol 74, 164–171, doi:10.1016/j.coi.2022.01.004 (2022).

21 Zhang, Z. et al. Multifaceted functions of STING in human health and disease: from molecular mechanism to targeted strategy. Signal Transduct Target Ther 7, 394, doi:10.1038/s41392-022-01252-z (2022).

22 Decout, A., Katz, J. D., Venkatraman, S. & Ablasser, A. The cGAS-STING pathway as a therapeutic target in inflammatory diseases. Nat Rev Immunol 21, 548–569, doi:10.1038/s41577-021-00524-z (2021).

23 Dixon, C. R. et al. STING nuclear partners contribute to innate immune signaling responses. iScience 24, 103055, doi:10.1016/j.isci.2021.103055 (2021).

24 Motwani, M., Pesiridis, S. & Fitzgerald, K. A. DNA sensing by the cGAS-STING pathway in health and disease. Nat Rev Genet 20, 657–674, doi:10.1038/s41576-019-0151-1 (2019).

25 Malik, P. et al. NET23/STING promotes chromatin compaction from the nuclear envelope. PLoS One 9, e111851, doi:10.1371/journal.pone.0111851 (2014).

26 Goldstone, D. C. et al. HIV-1 restriction factor SAMHD1 is a deoxynucleoside triphosphate triphosphohydrolase. Nature 480, 379–382, doi:10.1038/nature10623 (2011).

27 Coquel, F. et al. SAMHD1 acts at stalled replication forks to prevent interferon induction. Nature 557, 57–61, doi:10.1038/s41586-018-0050-1 (2018).

28 Graziano, S., Kreienkamp, R., Coll-Bonfill, N. & Gonzalo, S. Causes and consequences of genomic instability in laminopathies: Replication stress and interferon response. Nucleus 9, 258–275, doi:10.1080/19491034.2018.1454168 (2018).

29 Coll-Bonfill, N. et al. Progerin induces a phenotypic switch in vascular smooth muscle cells and triggers replication stress and an aging-associated secretory signature. Geroscience 45, 965–982, doi:10.1007/s11357-022-00694-1 (2023).

30 Chatzidoukaki, O. et al. R-loops trigger the release of cytoplasmic ssDNAs leading to chronic inflammation upon DNA damage. Sci Adv 7, eabj5769, doi:10.1126/sciadv.abj5769 (2021).

31 Lazarchuk, P., Nguyen, V. N., Brunon, S., Pavlova, M. N. & Sidorova, J. M. Innate immunity mediator STING modulates nascent DNA metabolism at stalled forks in human cells. Front Mol Biosci 9, 1048726, doi:10.3389/fmolb.2022.1048726 (2022).

32 Kreienkamp, R. et al. Vitamin D receptor signaling improves Hutchinson-Gilford progeria syndrome cellular phenotypes. Oncotarget 7, 30018–30031, doi:10.18632/oncotarget.9065 (2016).

33 Li, S. et al. Cytosolic DNA sensing by cGAS/STING promotes TRPV2-mediated Ca(2+) release to protect stressed replication forks. Mol Cell 83, 556–573 e557, doi:10.1016/j.molcel.2022.12.034 (2023).

34 Roy, S., Luzwick, J. W. & Schlacher, K. SIRF: Quantitative in situ analysis of protein interactions at DNA replication forks. J Cell Biol 217, 1521–1536, doi:10.1083/jcb.201709121 (2018).

35 Graziano, S. et al. Lamin A/C recruits ssDNA protective proteins RPA and RAD51 to stalled replication forks to maintain fork stability. J Biol Chem 297, 101301, doi:10.1016/j.jbc.2021.101301 (2021).

36 Kychygina, A. et al. Progerin impairs 3D genome organization and induces fragile telomeres by limiting the dNTP pools. Sci Rep 11, 13195, doi:10.1038/s41598-021-92631-z (2021).

37 Zhang, B., Xu, P. & Ablasser, A. Regulation of the cGAS-STING Pathway. Annu Rev Immunol 43, 667–692, doi:10.1146/annurev-immunol-101721-032910 (2025).

38 Hopfner, K. P. & Hornung, V. Molecular mechanisms and cellular functions of cGAS-STING signalling. Nat Rev Mol Cell Biol 21, 501–521, doi:10.1038/s41580-020-0244-x (2020).

39 Dunphy, G. et al. Non-canonical Activation of the DNA Sensing Adaptor STING by ATM and IFI16 Mediates NF-kappaB Signaling after Nuclear DNA Damage. Mol Cell 71, 745–760 e745, doi:10.1016/j.molcel.2018.07.034 (2018).

40 Guey, B. et al. BAF restricts cGAS on nuclear DNA to prevent innate immune activation. Science 369, 823–828, doi:10.1126/science.aaw6421 (2020).

41 Wardlaw, C. P. & Petrini, J. H. J. ISG15 conjugation to proteins on nascent DNA mitigates DNA replication stress. Nat Commun 13, 5971, doi:10.1038/s41467-022-33535-y (2022).

42 Raso, M. C. et al. Interferon-stimulated gene 15 accelerates replication fork progression inducing chromosomal breakage. J Cell Biol 219, doi:10.1083/jcb.202002175 (2020).

43 Green, A. M. et al. APOBEC3A damages the cellular genome during DNA replication. Cell Cycle 15, 998–1008, doi:10.1080/15384101.2016.1152426 (2016).

44 Situ, K. et al. BRCA2 deficiency and replication stress drive APOBEC3-Mediated genomic instability. Nat Commun 16, 9544, doi:10.1038/s41467-025-64578-6 (2025).

45 Xia, T., Konno, H., Ahn, J. & Barber, G. N. Deregulation of STING Signaling in Colorectal Carcinoma Constrains DNA Damage Responses and Correlates With Tumorigenesis. Cell Rep 14, 282–297, doi:10.1016/j.celrep.2015.12.029 (2016).

46 Su, X., Lu, W. & Rabinowitz, J. D. Metabolite Spectral Accuracy on Orbitraps. Anal Chem 89, 5940–5948, doi:10.1021/acs.analchem.7b00396 (2017).

47 Gonzalez-Suarez, I., Redwood, A. B. & Gonzalo, S. Loss of A-type lamins and genomic instability. Cell Cycle 8, 3860–3865, doi:10.4161/cc.8.23.10092 (2009).

48 Gonzalez-Suarez, I. et al. Novel roles for A-type lamins in telomere biology and the DNA damage response pathway. EMBO J 28, 2414–2427, doi:10.1038/emboj.2009.196 (2009).

