## Supplemental material for "STING causes replication stress and nascent DNA degradation via SAMHD1"

**THIS FILE INCLUDES:**

Supplemental Figure 1

Supplemental Materials and Methods: Tables 1 to 2

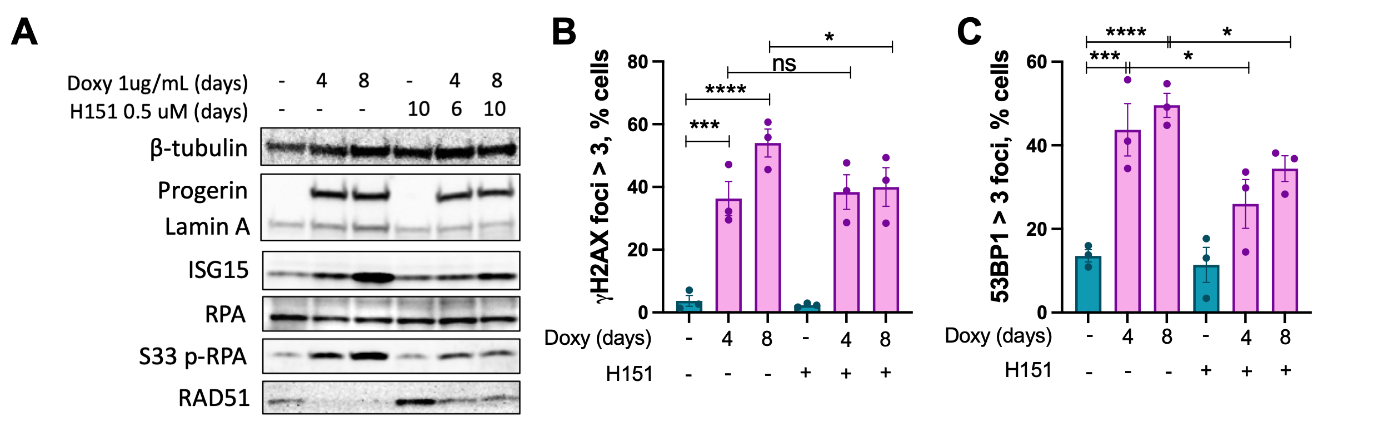

**Supplementary Figure 1. STING inhibition partially restores RAD51 expression and modestly reduces DNA damage markers in progerin-expressing cells.** (A) Immunoblot analysis of ^S33^p-RPA, RPA, and RAD51 levels in control HDF and progerin-expressing HDF that are treated with vehicle or STING inhibitor H151. β-tubulin was used as loading control. (**B**) IF staining with γH2AX antibody in HDF treated with doxycycline ± H151 for 4 or 8 days. Graph shows quantification of percentage of cells with ≥3 γH2AX foci per nucleus. (**C**) IF staining with 53BP1 antibody in HDF treated with doxycycline ± H151 for 4 or 8 days. Graph shows quantification of percentage of cells with ≥3 53BP1 foci per nucleus.

**Supplemental information about materials:**

**Table S1.** Antibodies and dilutions in this study.

| **Western blot** | | | |
| --- | --- | --- | --- |
| **Antibody** | **Dilution** | **Company** | **Catalog number** |
| β-tubulin | 1:5,000 | Origene | AP31823PU-N |
| Lamin A | 1:3,000 | Abcam | 26300 |
| Progerin | 1:1,000 | Santa Cruz | 81611 |
| H3 | 1:60,000 | Abcam | 1791 |
| ISG15 | 1:1,000 | Santa Cruz | 166755 |
| S172 p-TBK1 | 1:1,000 | Cell Signaling | 5483S |
| STAT1 | 1:1,000 | Cell Signaling | 14994S |
| S727 p-STAT1 | 1:1,000 | Cell Signaling | 9177S |
| STING | 1:1,000 | Cell Signaling | 13647 |
| S366 p-STING | 1:1,000 | Cell Signaling | 50907 |
| S386 p-IRF3 | 1:1000 | Cell Signaling | 37829 |
| RIG-I | 1:1,000 | Cell Signaling | 3743S |
| SEC61 | 1:1,000 | Cell Signaling | 14868 |
| SAMHD1 | 1:1000 | Cell Signaling | 49158S |
| GAPDH | 1:1,000 | Cell Signaling | 2118 |
| RPA | 1:1000 | Abcam | ab2175 |
| S33 p-RPA | 1:1000 | Bethyl | A300-246A |
| **Immunofluorescence** |  |  |  |
| STING | 1:500 | Cell Signaling | 13647 |
| S366 p-STING | 1:1200 | Cell Signaling | 50907 |
| 53BP1 | 1:1,000 | Santa Cruz | 22760 |
| γH2AX | 1:1,000 | Cell Signaling | 2577 |
| S33 p-RPA | 1:1000 | Bethyl | A300-246A |
| **PLA** |  |  |  |
| Lamin A | 1:500 | Abcam | AB26300 |
| STING | 1:100 | Invitrogen | MA5-26030 |
| Biotin | 1:200 | Cell Signaling | 5597S |

**Table S2.** Oligo sequences used in the study.

Dharmacon Catalog Item D-001810-10-20 ON-TARGETplus Non-targeting Pool: UGGUUUACAUGUCGACUAA, UGGUUUACAUGUUGUGUGA, UGGUUUACAUGUUUUCUGA, UGGUUUACAUGUUUUCCUA

Dharmacon Catalog Item L-024333-00-0005, ON-TARGETplus Human TMEM173/STING (340061) siRNA - SMARTpool, 5 nmol

ON-TARGETplus SMARTpool siRNA J-024333-05, TMEM173 Target Sequence: GGUCAUAUUACAUCGGAUA

ON-TARGETplus SMARTpool siRNA J-024333-06, TMEM173 Target Sequence: AACAUUCGCUUCCUGGAUA

ON-TARGETplus SMARTpool siRNA J-024333-07, TMEM173 Target Sequence: GCAUCAAGGAUCGGGUUUA

ON-TARGETplus SMARTpool siRNA J-024333-08, TMEM173 Target Sequence: GCACCUGUGUCCUGGAGU
